# Time-of-day-dependent responses of cyanobacterial cellular viability against oxidative stress

**DOI:** 10.1101/851774

**Authors:** Kenya Tanaka, Ginga Shimakawa, Shuji Nakanishi

## Abstract

As an adaptation to periodic fluctuations of environmental light, photosynthetic organisms have evolved a circadian clock. Control by the circadian clock of many cellular physiological functions, including antioxidant enzymes, metabolism and the cell cycle, has attracted attention in the context of oxidative stress tolerance. However, since each physiological function works in an integrated manner to deal with oxidative stress, whether or not cell responses to oxidative stress are under circadian control remains an open question. In fact, circadian rhythms of oxidative stress tolerance have not yet been experimentally demonstrated. In the present work, we applied an assay using methyl viologen (MV), which generates reactive oxygen species (ROS) under light irradiation, and experimentally verified the circadian rhythms of oxidative stress tolerance in photosynthetic cells of the cyanobacterium *Synechococcus elongatus* PCC7942, a standard model species for investigation of the circadian clock. Here, we report that ROS generated by MV treatment causes damage to stroma components and not to the photosynthetic electron transportation chain, leading to reduced cell viability. The degree of decrease in cell viability was dependent on the subjective time at which oxidative stress was applied. Thus, oxidative stress tolerance was shown to exhibit circadian rhythms. In addition, the rhythmic pattern of oxidative stress tolerance disappeared in mutant cells lacking the essential clock genes. Notably, ROS levels changed periodically, independent of the MV treatment. Thus, we demonstrate for the first time that in cyanobacterial cells, oxidative stress tolerance shows circadian oscillation.

## Introduction

All oxygenic photosynthetic organisms have evolved circadian clocks to adapt to the predictable, 24 h changes in light levels. Since daily fluctuations in environmental light levels lead to periodic variations in reactive oxygen species (ROS) levels in photosynthetic organisms, the physiological function of the circadian clock has been thought to be relevant to oxidative stress tolerance. For example, the activity or redox state of antioxidant enzymes are regulated by the circadian clock^1,2^. It has also been reported that circadian regulation of energy metabolism keeps ROS levels low^3^. Moreover, it has been suggested that circadian gating of cell division at a suitable time protects cells from photosynthetic oxidative stress^4^.

As mentioned above, the relationship between the circadian clock and oxidative stress has been described in studies of individual functions. However, since each adaptive function works in concert in natural environments, the effects of the circadian clock on cellular oxidative stress tolerance cannot be determined by analyzing each function. Although circadian control is important for cell physiology, the impact of circadian control of physiological systems on cells has not yet been fully elucidated.

The cyanobacterium *Synechococcus elongatus* PCC7942 (hereafter, *S. elongatus*), a photosynthetic autotrophic bacterium, is an appropriate model for study of circadian control of cellular physiology because the molecular mechanisms underlying its circadian clock is well understood, as summarized in recent reviews^5,6^.Although the circadian clock is known to regulate the expression patterns of many genes, this regulation is not necessarily reflected in protein abundance^7,8^. Moreover, to date, no phenotype showing circadian rhythm in oxidative stress tolerance has been experimentally confirmed. In this work, we developed an assay to examine the effect of oxidative stress on cell viability in an attempt to characterize the circadian rhythms of oxidative stress tolerance.

Generally, in photosynthetic living cells, the generation of large amounts of ROS is triggered by unfavorable conditions such as strong light irradiation or exposure to high (or low) temperature. However, these intense stimuli may mask the circadian control of oxidative stress tolerance because they directly affect a variety of metabolic pathways and induce adaptation mechanisms against oxidative stress, such as structural optimization and modification of light-energy conversion systems. A lack of detailed understanding of how ROS levels change in harsh conditions might hinder our overall understanding of oxidative stress tolerance. Therefore, we used methyl viologen (MV) with light irradiation as a ROS generation system with the aim of evaluating circadian clock involvement in oxidative stress tolerance control at the phenotype level.

## Results

### ROS generation profile

The molecular mechanism of ROS generation by MV has been well clarified and is well suited for the present work. Specifically, under light irradiation, MV mediates an electron transfer reaction from photosystem I (PSI) or ferredoxin to oxygen, resulting in the production of superoxide^9,10^. The generated superoxide is converted to H_2_O_2_ and a hydroxyl radical by a subsequent disproportionation reaction and Fenton type reaction^11^. In our study, *S. elongatus* cells grown in a turbidostat were assayed using a protocol in which 50 µM MV was added to cell culture aliquots, followed by 30 min light irradiation (this protocol for ROS generation is hereafter denoted “MV/light-treatment”). After MV/light-treatment, cells were washed with BG-11 medium and oxidative damage was evaluated by measuring oxygen evolution activity, P700 absorbance change or colony forming units (CFU) (Fig. 1).

**Figure 1.**
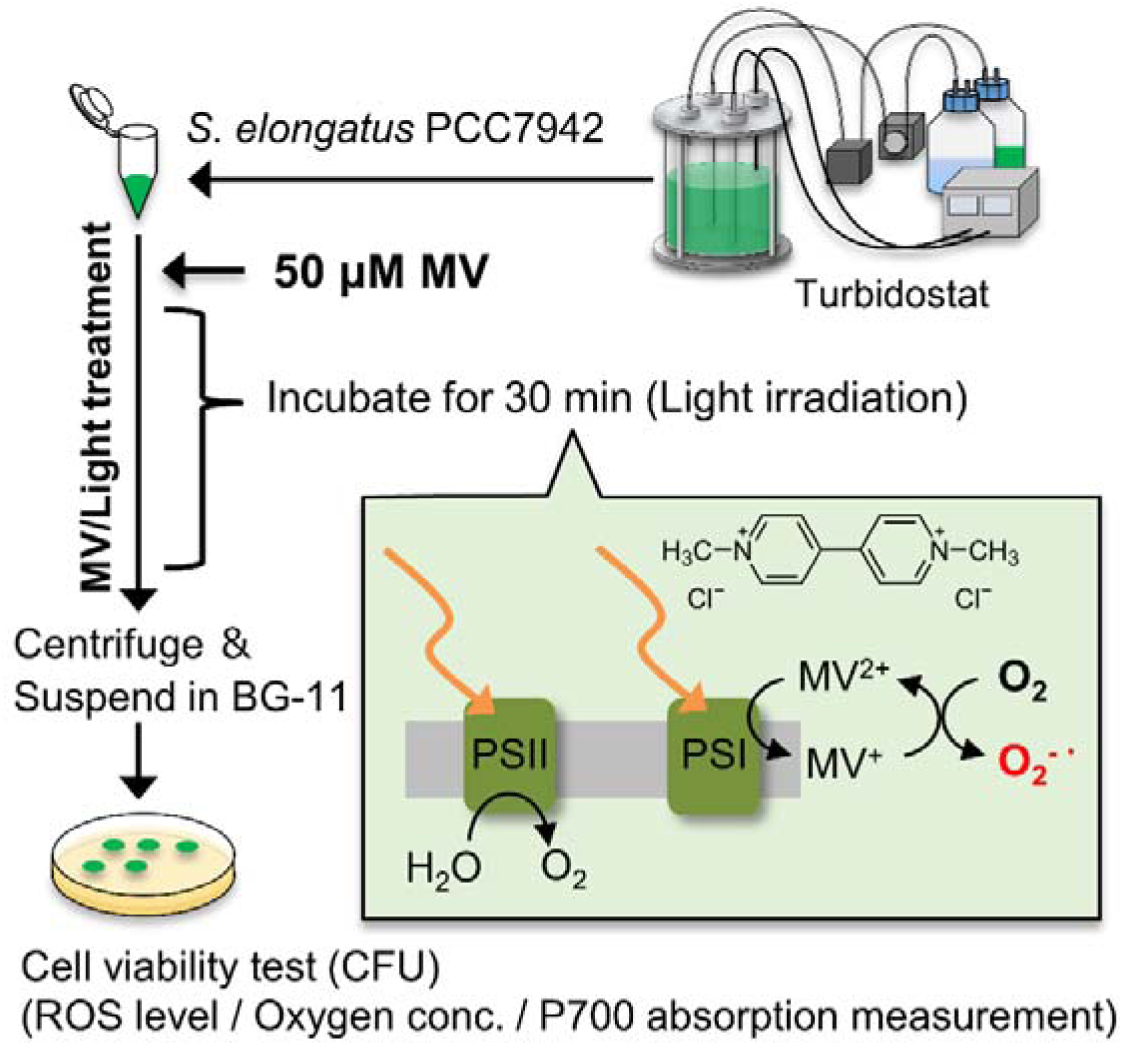
Schematic of *S. elongatus* growth, followed by ROS generation upon addition of MV with light incubation (MV/light treatment) and ROS tolerance evaluation. MV: methyl viologen, PSI: photosystem I, PSII: photosystem II.

ROS generation by MV/light treatment was confirmed by assay using 2’,7’-dichlorodihydrofluorescein diacetate (H_2_DCFDA), which is converted by intracellular esterases to 2’,7’-dichlorodihydrofluorescein, which can subsequently be oxidized by hydrogen peroxide, hydroxyl and peroxyl radical to a fluorescent compound with an emission peak at around 520 nm. Thus, intracellular ROS levels can be quantified by fluorescence intensity (Fig. 2A). To confirm light intensity dependency of ROS levels without the influence of the circadian clock, cyanobacterial cells were not entrained by LD cycles. The difference in fluorescence intensity was normalized to control samples incubated without MV in darkness, and showed a dependence on light intensity. A small increase in ROS levels was observed even for samples incubated in darkness with 50 µM MV. We attribute this small increase to the reduction of MV by intracellular NADPH.

**Figure 2.**
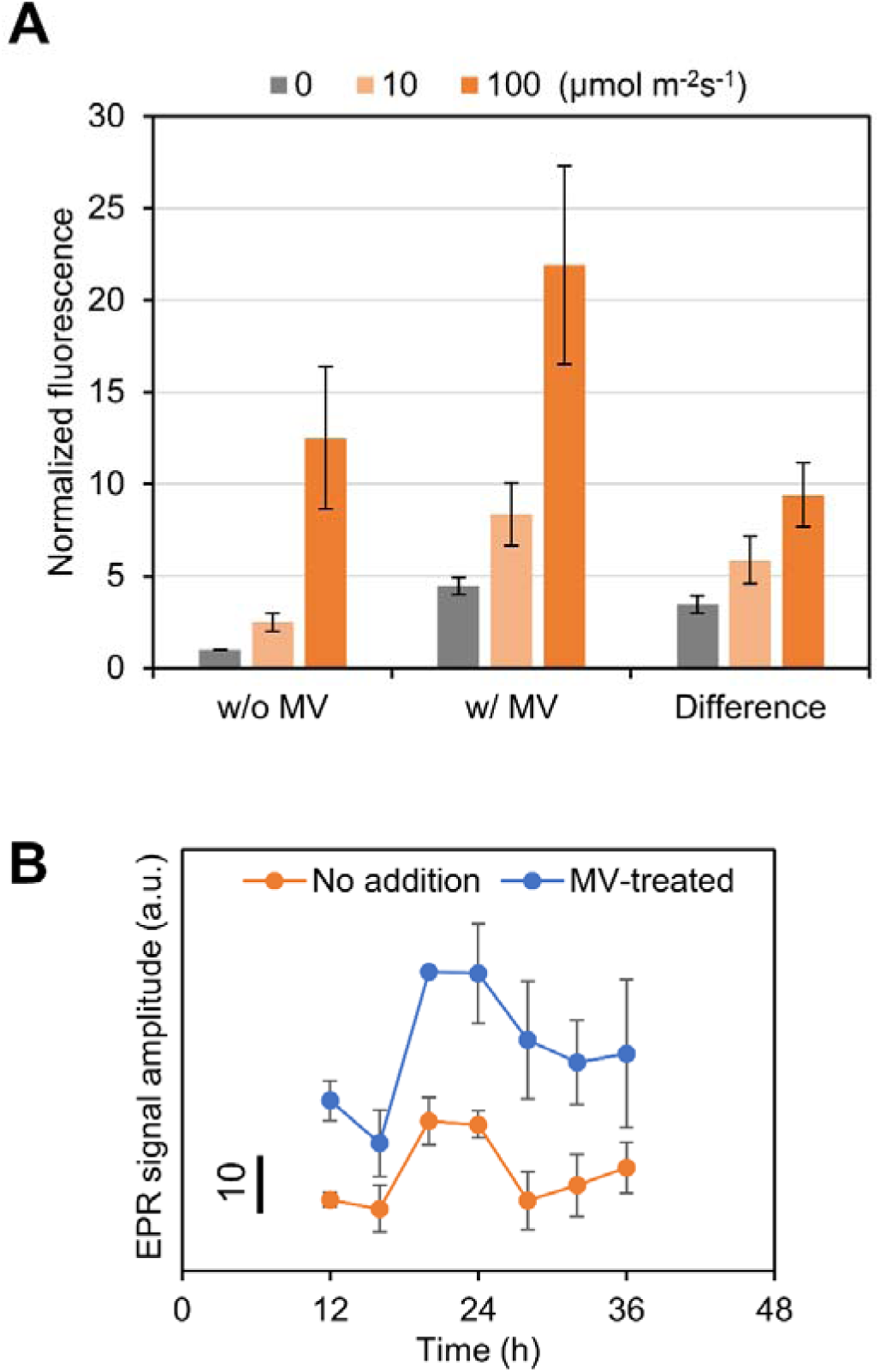
Effects of addition of MV, light irradiation intensity, and subjective time on intracellular ROS levels. (A) Wild type cells were incubated without MV (w/o MV) or with 50 µM MV (w/ MV) under three levels of light intensity (0, 10, and 100 µmol m^-2^ s^-1^) for 30 min. Fluorescence intensity of control samples (w/o H_2_DCFDA) was subtracted from the fluorescence intensity of samples with H_2_DCFDA, and the differences in fluorescence between with and without MV at each light intensity level are presented. Fluorescence intensity was measured from three biological replicates in each experiment. Values are means ± SD (bars) results from three independent experiments (therefore, each bar is the average of n = 9). (B) Dependency of ROS levels in wild type cells on circadian time. Wild type cells sampled at various subjective times were treated with 10 µmol m^-2^ s^-1^ light irradiation for 30 min, with or without 50 µM MV. Generated hydroxyl radicals were determined by spin trapping EPR spectroscopy with 4-POBN. Values are means ± SD (bars) of three biological replicates.

Although the assay using H_2_DCFDA is an easy method for evaluating ROS level, samples containing H_2_DCFDA cannot be cryopreserved due to their instability. Therefore, in order to confirm whether ROS level depends on the subjective time with H_2_DCFDA, it is necessary to perform the sampling and measurement at once at each time point. To overcome this technical difficulty, electron paramagnetic resonance (EPR) method was adopted to measure the time variation of ROS level. With this technique, a sample containing the spin-trapping agent can be frozen and stored, allowing us to collect samples over time and then conduct high-throughput experiments. Specifically, in cell suspensions subjected to MV/light treatment at various subjective times, levels of ROS, including superoxide radicals, hydrogen peroxide, and hydroxyl radicals, were determined by EPR using 4-pyridyl-1-oxide-N-tert-butylnitrone as a spin-trapping agent. We observed circadian rhythms in ROS levels in both samples treated and non-treated by MV/light-treatment with light intensity of 10 µmol m^-2^s^-1^ (Fig. 2B). Notably, we cannot make a quantitative discussion based on the absolute value of the EPR signal because baseline levels cannot be controlled. However, the EPR signal clearly exhibited circadian rhythms. Furthermore, consistent with the results of the H_2_DCFDA assay (Fig. 2A), MV/light treatment increased the average EPR signal value (Fig. S1).

### Effect of MV-induced oxidative stress on cell viability

Next, we examined the influence of MV/light treatment on cell viability. Treated cells were cultured on agar plates, and the number of colonies after 6 or 7 days was counted (Fig. 3A). To eliminate the effect of the circadian clock, cells that were not entrained to the LD cycle were used. Although ROS levels increased when cells were incubated in darkness with MV (Fig. 2A), there were no significant differences in colony-forming units (CFU) between samples incubated in weak light irradiation without MV versus darkness with MV. In contrast, for MV/light treated cells, CFU values decreased in a light intensity-dependent manner, indicating that ROS generated via the photosynthetic electron transport chain (PETC) through MV led to reduced cell viability. In addition, the finding that CFU changed in a light intensity-dependent manner during MV/light treatment means that CFU numbers were not influenced by residual MV during culture on agar plates, confirming that washing and serial dilution done prior to cell plating is sufficient to almost completely remove MV.

**Figure 3.**
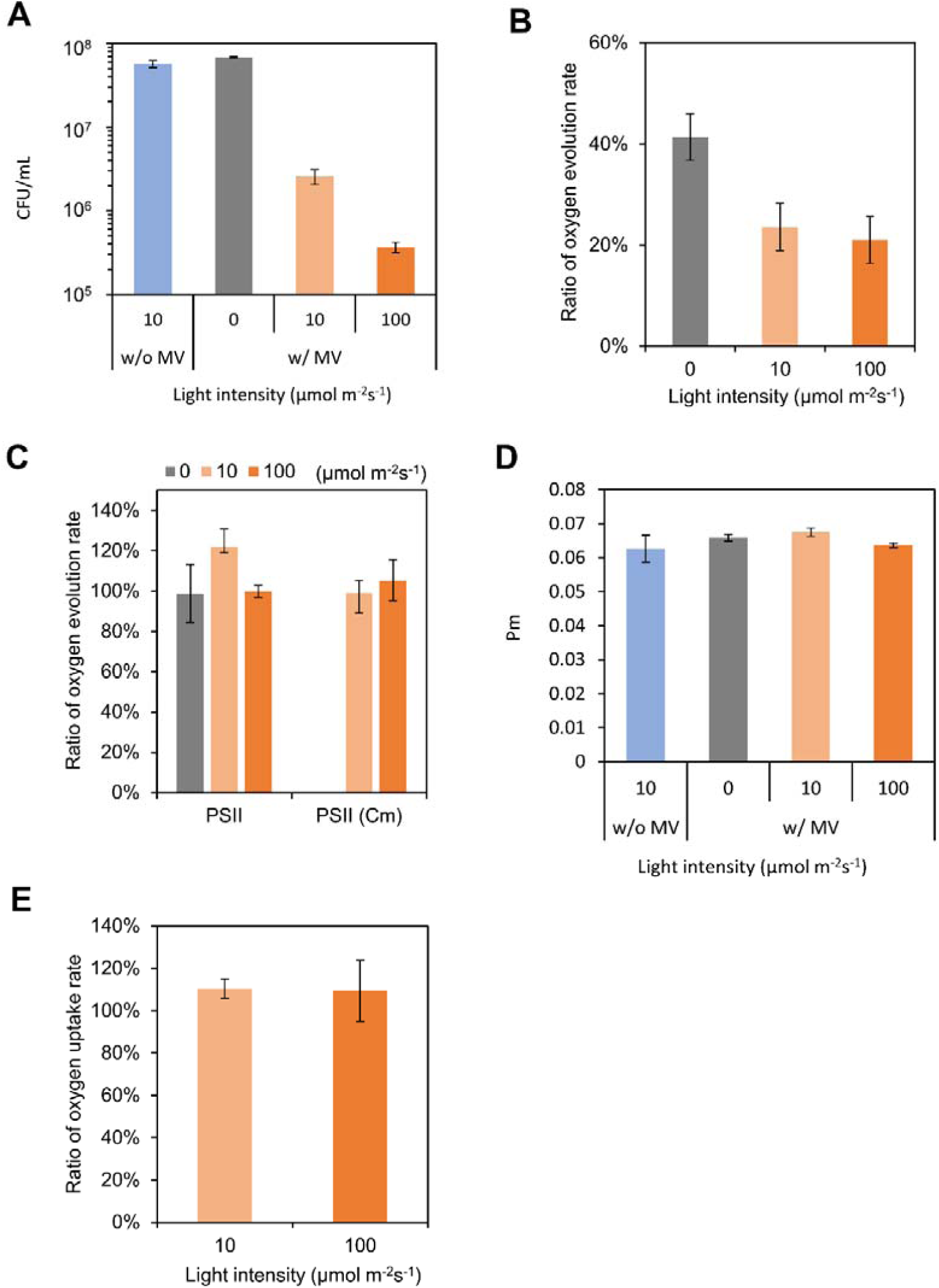
Evaluation of oxidative stress generated by MV/light treatment. (A) MV/light-treated cells were spotted on agar plates and grown to visualize the effects of MV/light treatment on cell viability. Obtained colonies were counted to calculate CFU. Values and error bars represent means ± SD of three technical replicates. The ratio of the oxygen evolution rate of 10 µg Chl/ml suspension-containing cells treated with and without 50 µM MV under three light intensities (MV/light treatment) was measured in the presence of (B) NaHCO3, (C) DCBQ, or DCBQ and 200 µg/ml chloramphenicol, indicated as PSII and PSII (Cm), respectively. (D) Oxidizable P700 (Pm). (E) Ratio of oxygen uptake rate of 10 µg Chl/ml suspension containing cells, treated with or without 50 µM MV to examine MV/light treatment-induced oxidative damage of the photosynthetic electron transport chain, including components from PSII to PSI. For measurement of oxygen uptake, 1 mM MV, 1 mM KCN and 10 mM methyl amine were added to a 10 µg Chl/ml suspension containing MV/light-treated cells in an oxygen electrode chamber. (A-D) Values and error bars represent means ± SD of three independent experiments.

### Effect of MV-induced oxidative stress on photosynthetic activity

To characterize oxidative stress damage by MV/light treatment in detail, we next investigated the effects of the treatment on the activities of overall electron transfer, including photosystem II (PSII) and photosystem I (PSI), activities. Overall electron transfer activity was evaluated by measuring the oxygen evolution reaction (OER) rate when CO_2_ (NaHCO_3_) was used as the sole electron acceptor. The ratio of the OER rate for cells incubated with MV versus that obtained for cells incubated without MV is shown in Fig. 3B and indicates that the OER rate markedly decreased following the addition of MV in the dark condition and decreased even more in higher light intensity conditions. These patterns are consistent with normalized fluorescence data (Fig. 2A), which reveal light intensity-dependent increase in ROS levels following the MV treatment in both dark and light conditions.

Next, PSII activity was evaluated by measuring the OER rate in the presence of 2,6-dichlorobenzoquinone (DCBQ), which intercepts photosynthetic electrons, mainly from the Q_B_ site in PSII. The OER rate was not affected by the addition of MV or light intensity (Fig. 3C) but was affected when CO_2_ was used as the electron acceptor (Fig. 3B). We propose that this is either because ROS generated by MV addition did not damage PSII or any damaged PSII was quickly repaired. To verify which of these two pathways led to the observed result, PSII activity measurements were next performed in the presence of a translation inhibitor, chloramphenicol (Cm), which prevents repair of PSII. Cells incubated in a medium containing Cm but not MV showed reduced PSII activity when exposed to light at an intensity of 100 µmol m^-2^s^-1^ (Fig. S2). However, PSII activity in the presence of Cm and at a light intensity of 10 µmol m^-2^s^-1^ was maintained at the same level as that observed in the absence of Cm. This indicates that photoinhibition of PSII does not occur under conditions of irradiation with 10 µmol m^-2^s^-1^. In contrast, MV/light treatment did not affect PSII activity, irrespective of light intensity (Fig. 3C). These results indicate that ROS generated under MV/light treatment does not damage PSII.

Oxidative damage to PSI was also evaluated using an established protocol based on the absorption spectroscopy for the reaction center of PSI (chlorophyll, P700)^12^. As shown in Fig. 3D, the amount of oxidizable P700 (Pm) in cells treated with MV was constant over the range of light intensity tested, indicating that MV/light treatment did not damage PSI.

Finally, the effect of MV-induced oxidative stress on PETC, including all components from PSII to PSI, was tested by measuring the light-induced O_2_ uptake rate, with H_2_O and MV serving as the electron donor and accepter, respectively. As shown in Fig. 3E, cells subjected to MV/light treatment maintained electron transfer activity at the same level for light irradiation at 10 and 100 µmol m^-2^s^-1^. Taken together, generated ROS did not damage PETC, which is itself the machinery for generating ROS.

Oxidative damage to PSII or PSI is known to be caused by ROS generation due to excessive reduction of PETC components but not by ROS generation due to external addition of H_2_O_2_ or MV^13-15^. In good agreement with these studies, as mentioned earlier, PETC activity was not impacted by oxidative stress induced by MV/light treatment. Nevertheless, overall photosynthetic activity and CFU values were decreased following MV/light treatment (Figs. 3A, B). These results clearly indicate that stromal components, including enzymes participating in electron flux downstream of PSI, were damaged by ROS. In fact, lipids, DNA and Rubisco, which has CO_2_ fixing ability, are known to be damaged by ROS generated by MV^16,17^. Oxidative damage to such components is likely to have led to a decrease in overall electron transfer activity and CFU.

### Effect of the circadian clock on MV-induced oxidative stress tolerance

To investigate circadian control of oxidative stress tolerance, MV/light treatment was used to generate ROS in cells sampled at various subjective times. For these tests, cells cultured in a turbidostat were kept at a constant cell density (optical density 730 nm (OD_730_) = 0.6 ± 0.1, Fig. S3) in order to exclude the possibility that differences in growth stage influence the CFU results. Light intensity during MV/light treatment was set to 10 µmol m^-2^s^-1^ to avoid photoinhibition of PSII (Fig. S2). After subjecting the cells to light-dark (LD) cycles with 24 h periodicity, cells were exposed to continuous light (LL) conditions in the turbidostat (Figs 4A and S4). While non-treated wild type (WT) cells (filled circles) showed no dependence on subjective time, MV/light-treated cells showed circadian rhythm (open circles). Mutant cells lacking clock genes (Δ*kaiABC*) did not show clear rhythms, whether or not they were treated with MV/light (closed and open squares in Fig. 4A). Importantly, the oscillatory phase of the CFU rhythm was determined by the phase of LD cycles (Fig. 4B). Further, using period mutant cells (F470Y, S157P) with a free running period (FRP) of 16 and 21 h, respectively^18^, CFU results showed a periodicity that corresponded with the FRP (Fig. 5).

**Figure 4.**
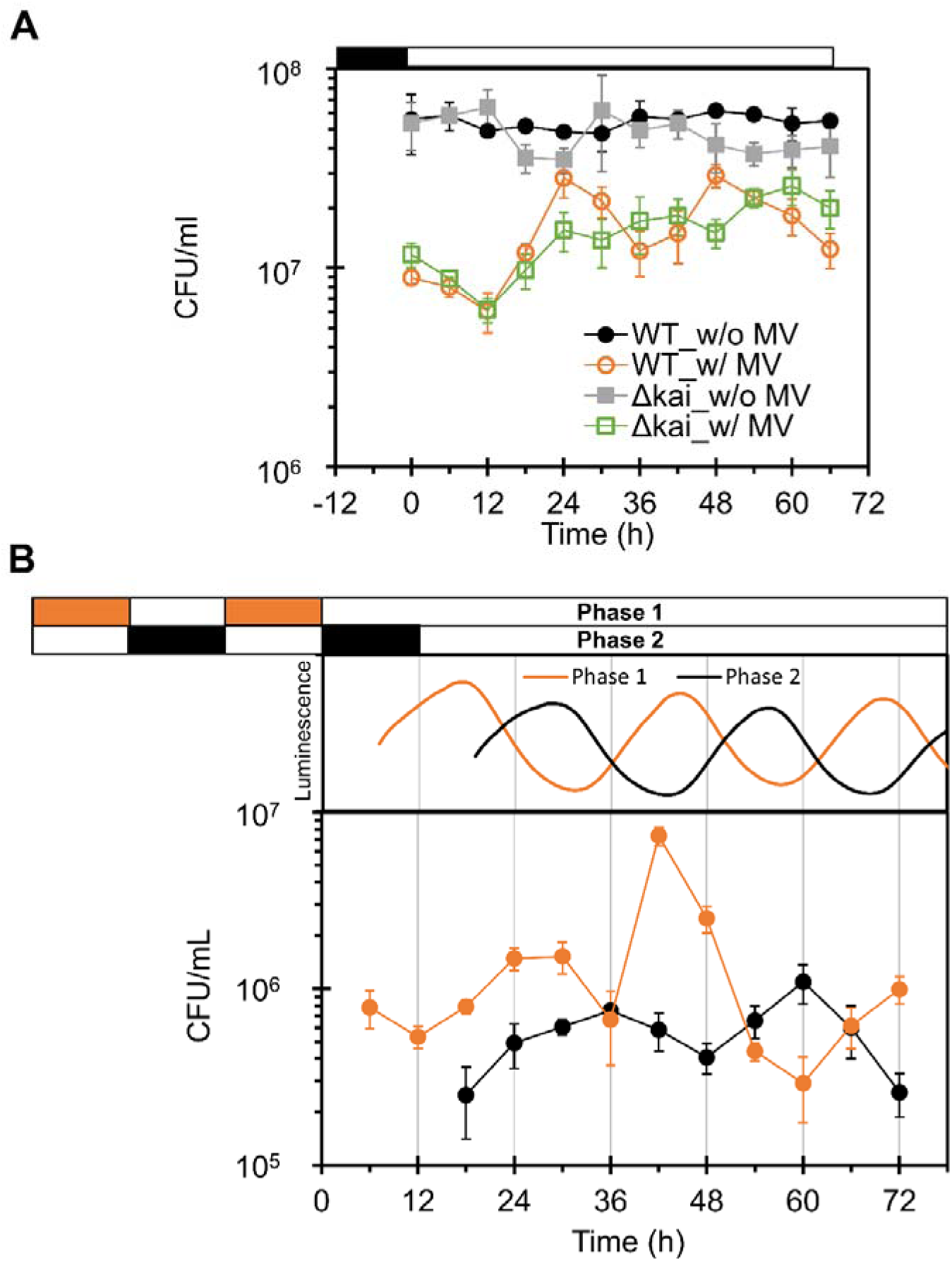
Time series of CFU of MV/light treated cells. (A) Wild type (WT) and *kaiABC*-deficient mutant (Δ*kaiABC*) cells were harvested from turbidostat every 6 h, followed by MV/light treatment (10 µmol m^-2^ s^-1^) of each aliquot. Oxidative stress tolerance was visualized as the number of colonies that grew and (B) ROS tolerance of WT cell cultures entrained in antiphase were tested every 6 h. Luminescence rhythms of WT cells were also measured to identify the phase of each culture, and are shown above the CFU graph. White and black or orange bars denote light-dark (LD) cycles. Values and error bars represent means ± SD of three technical replicates.

**Figure 5.**
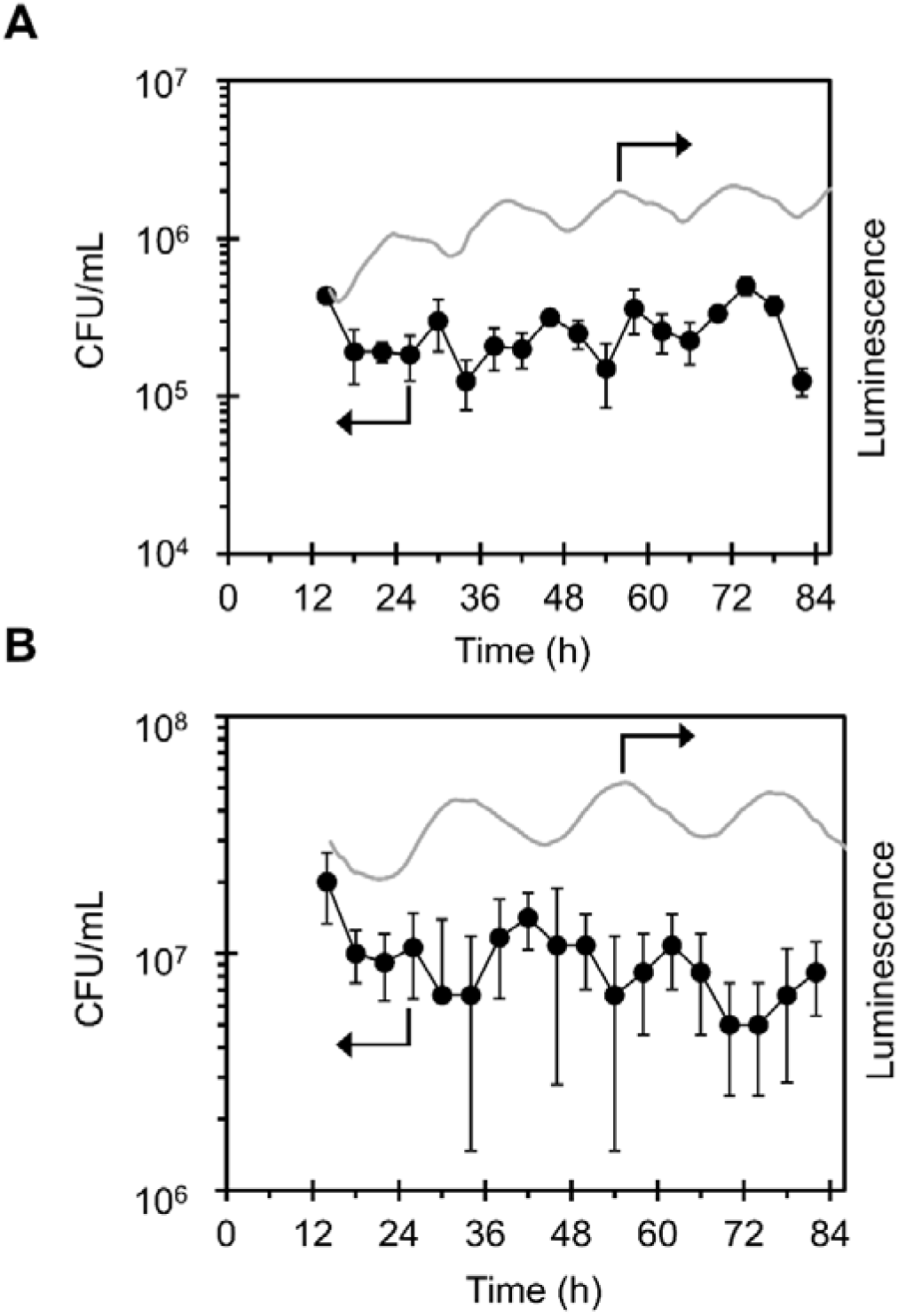
Correlation of period length between gene expression rhythms and ROS tolerance rhythms of period mutant strains. ROS tolerance of (A) F470Y and (B) S157P strains cultured in continuous light conditions following 12 h light/12 h dark cycles for entrainment were tested. Gray lines indicate luminescence traces of the cells, which confirm the expected short-period phenotype. Values represent means of three technical replicates.

## Discussion

As shown above, the circadian rhythms of oxidative stress tolerance were experimentally observed by using MV/light treatment, which causes oxidative damage in stromal components (Figs. 2-4). In the *kaiABC* deletion mutant, the CFU rhythm disappeared (Fig. 4A). In addition, short-period mutants showed CFU rhythms corresponding to each FRP (Fig. 5). These results clearly indicate that ROS management of the KaiABC system affects cell viability in a time-of-day-dependent manner. The time-of-day-dependent change in cell viability can arise from rhythmic variation in ROS levels (case-I) and/or by rhythmic damage of a critical component for cell viability (case-II), the latter of which can occur even at a constant ROS level. Considering that the circadian clock based on the KaiABC system is a global regulator, the effects of both of these factors are likely to change in a circadian manner.

First, let us consider case-I. As shown Fig. 2B, the ROS level of samples in natural conditions (i.e. without MV/light treatment) also varied in a subjective time-of-day-dependent manner. ROS levels are defined by the balance between the amount of ROS generated and the amount scavenged. In general, accumulation of ROS occurs when electron acceptor capacity at the PSI side is not high enough as compared with electron flux from the donor side^14^. Independent transcriptome studies showed that the expression patterns of genes encoding photosynthetic components related to the electron acceptor capacity at PSI, such as genes encoding Rubisco and GAPDH, are rhythmic under the control of KaiABC^7,19^. Thus, it is highly likely that the amount of ROS generated oscillates in a circadian manner, even in LL conditions. On the other hand, generated superoxide and H_2_O_2_ should be scavenged by antioxidant enzymes such as superoxide dismutase (SOD) and catalase. According to the microarray analyses, genes encoding of 2-cys peroxiredoxin, SOD and catalase are rhythmically expressed even in a LL condition^7,19^. These facts suggest that circadian variation in ROS levels is due at least in part to rhythmic gene expression as controlled by KaiABC.

Importantly, although circadian fluctuations in ROS levels were observed in both the presence and absence of MV/light treatment (Fig. 2B), circadian fluctuations in CFU were observed only for MV/light-treated cells (Fig. 4A). This inconsistence led us to hypothesize that cell viability changes only when ROS levels exceed a certain threshold. In other words, without MV/light treatment, the amount of ROS is not high enough to cause a decrease in CFU even at peak levels and thus, time-of-day-dependent changes in CFU were not observed. When the cells were MV/light-treated, average ROS levels were increased at all time points (Fig. 2B). However, ROS levels exceeded the threshold in subjective midnight, leading to circadian variation in cell viability. Consistent with this hypothesis, despite the fact that the addition of MV in dark increased ROS levels (Fig. 2A) and decreased photosynthetic activity (Fig. 3B), CFU values in this case showed no differences from those obtained in the absence of MV (Fig. 3A). The phases of circadian rhythms of ROS levels and CFU variation did not coincide: ROS levels peaked during a period from subjective midnight to dawn, whereas CFU peaked around subjective dawn. This could be because the ROS-induced response continues for several hours after MV/light treatment.

We also considered the possibility that rhythmic damage to a critical component required for cell viability (i.e., case-II) is the origin of the time-of-day-dependent variation. For example, it is known that cell division is controlled by circadian clock in order to help cope with oxidative stress^4^. Furthermore, in *S. elongatus*, the circadian clock regulates gating of cell division^20^. Considering these reports, we reason that circadian rhythm in CFU might have appeared due to the influence of circadian gating of cell division. We speculate that both of these two cases work in parallel and under the regulation of KaiABC, together leading to time-of-day-dependent changes in CFU that we observed.

In summary, this study experimentally demonstrated for the first time that ROS levels and cell viability in response to oxidative stress change in a circadian manner under the control of KaiABC. Circadian variation in CFU in response to MV/light treatment was observed in specific conditions (MV concentration: 50 µM, light intensity: 10 µmol m^-2^s^-1^, and light irradiation: 30 min). In principle, rhythmicity in CFU might also be observed, for example, at higher light intensity if the MV concentration was reduced or no MV was added. An important finding obtained throughout this work is that the intracellular ROS levels (case-I) and/or ROS sensitivity of a specific cellular component (case-II) can show circadian oscillation. We anticipate that further studies along this line will contribute to understanding competitive fitness advantages, which have attracted attention from the viewpoint of understanding the origin and evolution of the circadian clock^1,5^.

## Methods

### Bacterial strains and cell culture conditions

We used the following *S. elongatus* strains: wild-type reporter strains (P_*kaiBC*_*::luxAB* and P_*psbAI*_*::luxCDE*)^21^, a *kaiABC*-deficient P_*kaiBC*_ reporter strain^22^ and two *kaiC* mutant strains with short periodicity carrying P_*kaiBC*_ reporters (F470Y and S157P)^18^. These strains were grown in liquid or on solid (1.5% Bacto agar) BG-11 medium. Liquid cultures were grown in flasks (batch culture) or turbidostat (continuous culture) at 30 °C with air bubbling and exposure to light at an intensity of 30-40 µmol m^-2^s^-1^. For continuous culture, cell density was continuously monitored using an optical sensor with near-infrared light (NI; 840 nm ∼ 910 nm) and cell density was controlled by dilution with fresh medium when the OD value in the NI region exceeded a certain value.

### Methyl viologen (MV) treatment

MV was added at a final concentration of 50 µM to liquid culture medium to achieve an optical density at 730 nm (OD_730_) of 0.65±0.05. Negative control samples without MV were prepared at the same time. The samples were incubated for 30 min at the specified light intensity, and cells were collected by 5-min centrifugation at 4,400 xg at 30 °C, re-suspended in fresh BG-11 medium to remove MV, centrifuged again, and re-suspended again in fresh BG-11 to produce the cell density required for subsequent measurements.

### H_2_DCFDA assay

Intracellular ROS levels were measured as described previously with slight modifications^3,23^. Just before the assay, H_2_DCFDA (Invitrogen) was dissolved in ethanol to produce a 15 mM stock solution. The H_2_DCFDA stock solution or ethanol (vehicle) was then added to a liquid culture of OD_730_ of 0.65±0.05 and the treated culture was incubated in darkness for 30 min at 30 °C. Next, H_2_DCFDA was removed from the culture medium by centrifugation, the culture medium was combined with an equal amount of fresh BG-11 medium, and 50 µM MV was or was not added. To generate ROS, the cell suspension was incubated for 30 min at the indicated light intensity at 30 °C. After light treatment, the cells were placed in a 96-well plate and the fluorescence emission at 535 nm was measured (485 nm excitation) on a Tecan Infinite M200 Plate Reader. The background fluorescence signal of samples without H_2_DCFDA (ethanol only) was subtracted from the fluorescence values of the corresponding samples with H_2_DCFDA.

### Spin trapping EPR spectroscopy

Spin trapping assays with the spin probe 4-pyridyl-1-oxide-N-tert-butylnitrone (4-POBN; Sigma-Aldrich, St. Louis, USA) to detect the formation of hydroxyl radicals^24^ were carried out using cyanobacterial cells (OD_730_ = 0.65) in 10 mM sodium phosphate buffer (pH 7.0) containing 50 mM 4-POBN, 4% (v/v) ethanol, 50 µM Fe-EDTA, and 50 µM MV. After a 30-min incubation in the light (10 µmol m^-2^ s^-1^), the samples were centrifuged at 7,000 *g* for 1 min, and the supernatants were frozen in liquid nitrogen and stored at −80 °C for electron paramagnetic resonance (EPR) spectra analysis. The EPR spectra were recorded at room temperature in a standard quartz flat cell using an ESP-300 X-band spectrometer (Bruker, Rheinstetten, Germany). The following parameters were used: microwave frequency, 9.73 GHz; modulation frequency, 100 kHz; modulation amplitude, 1 G; microwave power, 6.3 milliwatt; receiver gain, 2 × 104; time constant, 40.96 ms; number of scans: 4.

### Measurement of oxygen concentration

Reaction mixtures containing BG-11 with 40 mM TES-NaOH (pH 7.5), 50 mM NaHCO_3_ and cells after MV/light treatment (10 µg Chl/mL) were stirred with a magnetic stirrer, and the oxygen concentration was monitored using an oxygen electrode (Hansatech, UK). Overall activity was determined by light irradiation with an LED source (pE-100^wht^, BioVision Technologies, USA) through a 550 nm long pass filter at light intensity of 660 µmol m^-2^s^-1^. Next, 0.4 mM 2,6-dichlorobenzoquinone (DCBQ) was added to determine PSII activity. To determine electron transfer activity from PSII to PSI, light-induced oxygen uptake of MV/light-treated cells was measured in the presence of 1 mM KCN, 1 mM MV, and 10 mM methyl amine.

### Photochemical measurement of total P700

Absorbance changes of P700 (reaction center chlorophyll of PSI) were monitored using a fiber version of the Dual-PAM-100 measuring system at room temperature. Samples containing BG-11 with 40 mM TES-NaOH (pH 7.5) and cells post-MV/light treatment (15 µg Chl/mL) were introduced into a cuvette. To measure maximum P700 absorption, saturating pulse illumination (10,000 µmol m^-2^s^-1^) was applied after irradiation with blue light (750 µmol m^-2^s^-1^) for 5 s.

### Viable cell plating

Samples after MV/light treatment were re-suspended in fresh BG-11 to reach an OD_730_ of 0.3. Each sample was serially diluted 1:10 four times in fresh BG-11 and then 4 µL of each diluted sample was spotted onto solid BG-11 plates without antibiotics. The plates were incubated at 30 °C under 30 µmol m^-2^s^-1^ constant light for 6 or 7 days and then photographed.

## Data availability

Data supporting the findings of this paper are available from the corresponding authors upon reasonable request.

## Acknowledgments

We thank Prof. K. Terauchi (Ritsumeikan University) and Prof. T. Kondo (Nagoya University) for kindly providing the mutant strains used in this work; and we thank Prof. K. Tanaka and Mr. T. Tsurumaki (Tokyo Institute of Technology) for providing a key suggestion regarding clock-controlled ROS tolerance through a presentation at the 59th Annual Meeting of the Japanese Society of Plant Physiologists (Sapporo, Japan 2018). This work was partially supported by the Advanced Low Carbon Technology Research and Development Program (JPMJAL1402) of the Japan Science and Technology Agency (JST), JSPS KAKENHI (19K22232 and 18J20176), and a JSPS oversea research fellowship (201860126).

## Author contributions

K.T. and S. N. designed the research. G.S. determined ROS levels by spin trapping EPR spectroscopy. K.T. performed all of the other experiments. K.T. designed the experiments. K.T. and S.N. wrote the first draft of the manuscript. K.T. prepared figures. All authors (K.T., G.S. and S.N.) edited the manuscript.

