## Supplemental material for "Time-of-day-dependent responses of cyanobacterial cellular viability against oxidative stress"

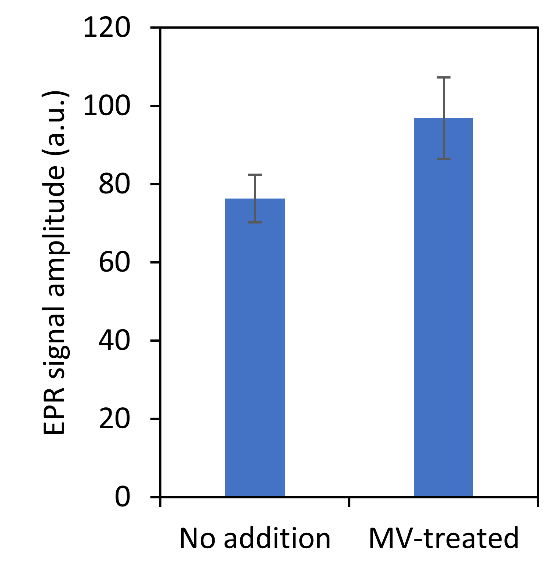

**Figure S1.** Effect of ROS level measured by spin trapping EPR spectroscopy on presence of 50 μM Methyl viologen (MV). The EPR signal values for different subjective times were averaged (Data is identical with Fig. 2B). Values are means ± SD of average values at seven different subjective time.

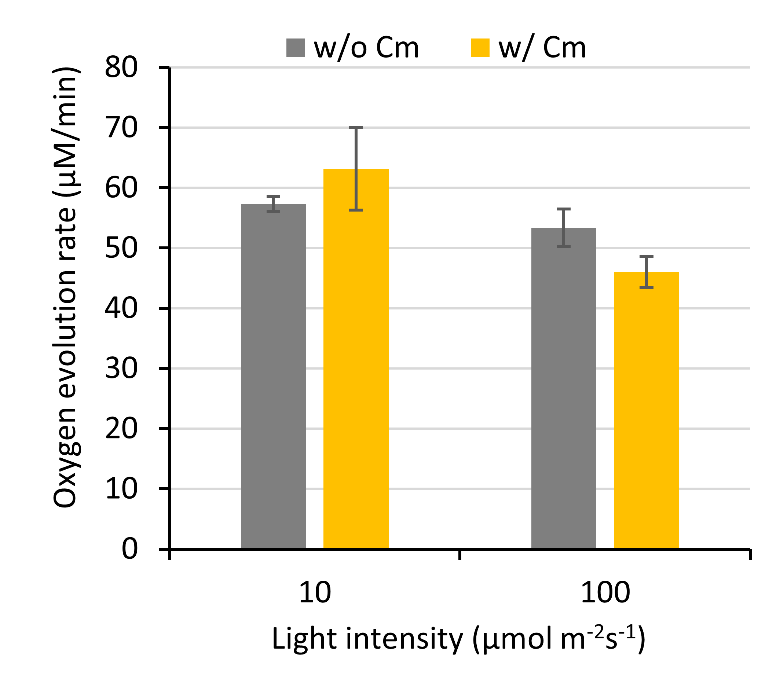

**Figure S2.** Effect of chloramphenicol (Cm) and light irradiation on PSII activity. Cells were incubated with or without 200 μg/mL Cm under light irradiation (10 or 100 µmol/m^2^/s) for 30 min. Oxygen evolution rate of reaction mixture containing BG-11 (40 mM TES-NaOH, pH 7.5), the treated cells (10 µg Chl/mL) and 0.4 mM 2,6-dichlorobenzoquinone (DCBQ) was measured under actinic light irradiation. Oxygen evolution rate of the cells incubated with Cm under 100 µmol/m^2^/s light irradiation was significantly lower than that of cells incubated without Cm (student t-test, p < 0.05), while cells incubated under 10 µmol/m^2^/s light irradiation showed no significant difference between with and without Cm. Values are means ± SD (bars) results from three independent experiments.

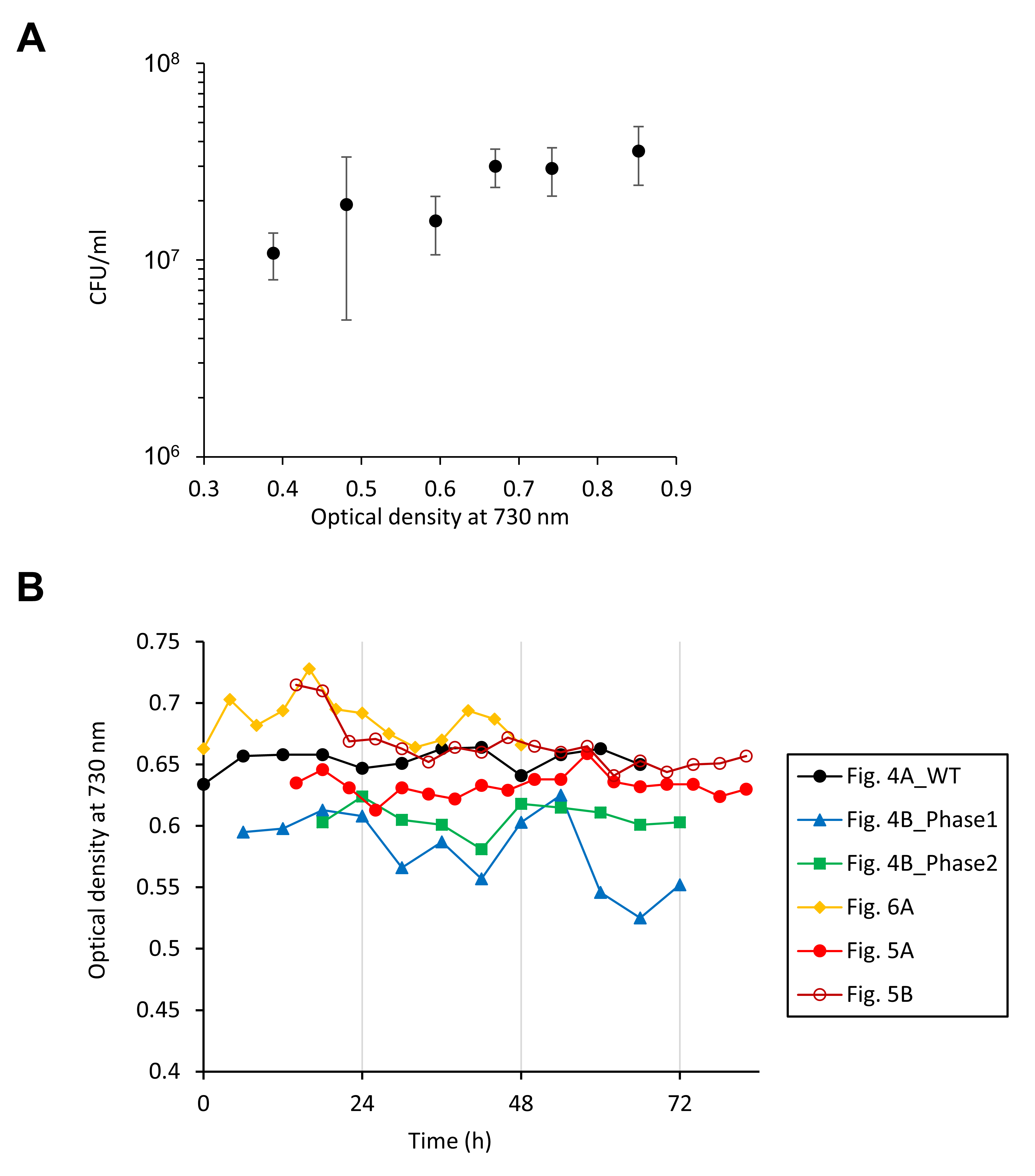

**Figure S3.** Effect of cell density during the MV/Light treatment on colony forming unit (CFU). (A) WT cell suspensions with various cell density (measured as optical density at 730 nm; OD_730_) were treated by the MV/Light treatment, followed by growing on agar plates. Values are means ± SD (bars) results from three biological replicates. (B) Time courses of OD_730_ values of cells taken for the experiments in which time-dependent ROS-tolerance change in continuous light condition was investigated (Figs. 4, 5, 6A). The time courses of the OD_730_ are hardly rhythmic, and the perturbations of the OD_730_ values are suggested not to cause significant CFU difference, indicating that the rhythmic CFU changes observed in Figs. 4, 5, 6A were not attributed to effect of time dependent cell density change.

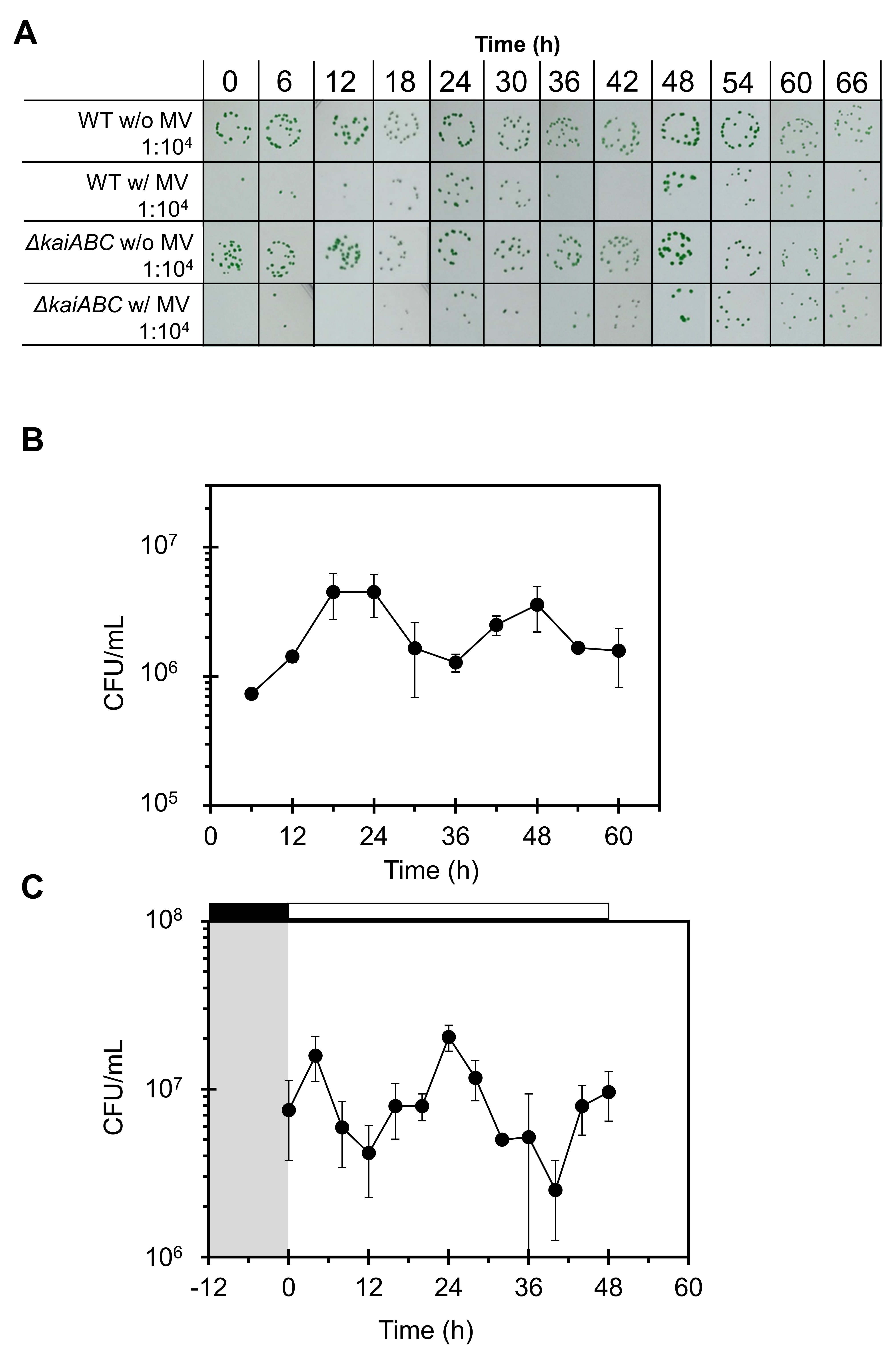

**Figure S4.** Time series of ROS-stress tolerance of cells grown in continuous light condition. (A) Representative photograph of colonies of samples in Fig. 4A are shown. (B, C) Circadian rhythm of ROS-tolerance tested with WT cell culture independent from that of Fig. 4. Values are means ± SD (bars) results from three technical replicates.
